# Pan cancer patterns of allelic imbalance from chromosomal aberrations in 33 tumor types

**DOI:** 10.1101/635193

**Authors:** Smruthy Sivakumar, F Anthony San Lucas, Yasminka A Jakubek, Jerry Fowler, Paul Scheet

**Affiliations:** Department of Epidemiology, The University of Texas MD Anderson Cancer Center, Houston, Texas, USA; The University of Texas MD Anderson Cancer Center UTHealth Graduate School of Biomedical Sciences, Houston, Texas, USA

**Keywords:** allelic imbalance, genomic instability, copy number alterations, cancer aneuploidy

## Abstract

Somatic copy number alterations (SCNAs), including deletions and duplications, serve as hallmarks of tumorigenesis. SCNAs may span entire chromosomes and typically result in deviations from an expected one-to-one ratio of alleles at heterozygous loci, leading to allelic imbalance (AI). The Cancer Genome Atlas (TCGA) reports SCNAs identified using a circular binary segmentation (CBS) algorithm, providing segment mean copy number estimates from Affymetrix single-nucleotide polymorphism DNA microarray total (log R ratio) intensities, but not allele-specific (“B allele”) intensities that inform of AI. Here we seek to provide a TCGA-wide description of AI in tumor genomes, including AI induced by SCNAs and copy-neutral loss-of-heterozygosity (cnLOH), using a powerful haplotype-based method applied to allele-specific intensities. We present AI summaries for all 33 tumor sites and propose an automated adjustment procedure to improve calibration of existing SCNA calls in TCGA for tumors with high levels of aneuploidy where baseline intensities were difficult to establish without annotation of AI. Overall, 94% of tumor samples exhibited AI. Recurrent events included deletions of 17p, 9q, 3p, amplifications of 8q, 1q, 7p as well as mixed event types on 8p and 13q. The AI-based approach identified frequent cnLOH on 17p across multiple tumor sites, with additional site-specific cnLOH patterns. Our findings support the exploration of additional methods for robust automated inference procedures and to aid empirical discoveries across TCGA.

## INTRODUCTION

Chromosomal instability resulting in somatic copy number alterations (SCNAs), including whole chromosomal or whole-arm gains and losses, impact large regions in the genome that might encompass known oncogenes and tumor suppressor genes. Such genomic instability events serve as hallmarks of tumorigenesis (Negrini et al. 2010) by resulting in the rapid accumulation of additional and possibly driver mutations. Previous studies have identified the presence of chromosomal SCNAs that are characteristic to specific tumor types and subtypes (Taylor et al. 2018; Ried et al. 2012; Hoadley et al. 2014). Consequences of chromosomal aberrations include altered responses to therapeutic regimens, gene expression modulations favoring increased cell proliferation and reduced expression of immune markers (Hutchinson 2017; van Jaarsveld and Kops 2016). Therefore, the accurate detection, genome-wide characterization and cataloging of such aberrations may help better elucidate their roles in tumor initiation or progression through correlative analyses with epidemiological variables or clinical outcomes.

The Cancer Genome Atlas (TCGA) provides a large repository of tumor specimens across multiple tissue types and varied clinicopathological features, thus facilitating several pan-cancer studies of cancer aneuploidy and tumor-specific copy-number signatures, such as those derived from single nucleotide polymorphism (SNP) genotyping array platforms (Taylor et al. 2018; Zack et al. 2013; Hoadley et al. 2014). However, studies of chromosomal SCNAs are challenging owing to the difficulty in acquiring high cellularity tumor specimens as well as limitations in automated computational approaches for the detection of these changes. However, unique challenges compound the automated detection of chromosome arm-level SCNAs from SNP arrays. First, the samples often derive from a mixture of cells from tumors and those which are “pathologically normal”, thereby necessitating algorithms with increased sensitivity to identify “subtle” genomic alterations in bulk samples of intermediate or modest cellularity. Second, most SCNA detection methods report genomic regions of copy number alterations and their segment mean copy number (CN) estimates, the characterization of which heavily relies on accurate identification of non-aberrant regions of the genome to establish a baseline signal intensity representative of neutral CN; however, tumor samples exhibiting high levels of genomic instability pose a challenge for such analyses.

To overcome these challenges, we sought to identify regions in the genome that exhibit allelic imbalance (AI), a deviation from the expected 1:1 ratio at germline heterozygous loci, a natural consequence of SCNAs such as duplications, deletions and copy-neutral loss-of-heterozygosity (cnLOH), using a haplotype-aware statistical method that is suitable for detecting megabase-scale aberrations (Vattathil and Scheet 2013). Outputs from SNP DNA microarrays include the following two measurements per marker: the “B allele” frequency (BAF), representing the proportion of the arbitrarily-labeled “B allele” at a locus; and the log R ratio (LRR), the total intensity of (both) allelic probes at the locus. In contrast to methods that utilize LRR to identify regions of SCNAs, hapLOH utilizes the BAF to identify regions of allelic imbalance, followed by a BAF and LRR threshold-based characterization of the identified events. In addition to being a more sensitive approach that identifies additional SCNAs (Figure 1A), a BAF-based method for AI detection also provides a relatively unexplored, yet informative, class of chromosomal aberrations -- cnLOH (Figure 1B). These represent regions of zero net copy number change but an extreme alteration in the ratio of alleles (i.e., change of germline heterozygous loci AB to AA or BB). The landscape of large cnLOH regions remains largely unknown due to the lack of sensitive algorithms for their automated detection. Recent pan-cancer investigations have identified evidence of focal LOH events accompanying mutations in genes involved in DNA damage repair pathways (Knijnenburg et al. 2018), as well as those accompanying polymorphisms in essential genes that result in cancer cell-specific vulnerabilities (Nichols et al.). However, very few studies have described the vital role of large, chromosome-arm level cnLOH in the development of hematologic malignancies (O’Keefe et al. 2010; Stirewalt et al. 2014; Schwartz and Papenhausen 2017), gastrointestinal tumors (Lourenço et al. 2014) and colorectal cancer (Melcher et al. 2011). Therefore, it is crucial to understand and identify the landscape of these cnLOH events across tumor sites to better understand their role in the complex mechanisms of tumorigenesis such as those contributing to the multi-hit pathogenesis of tumors.

**Figure 1.**
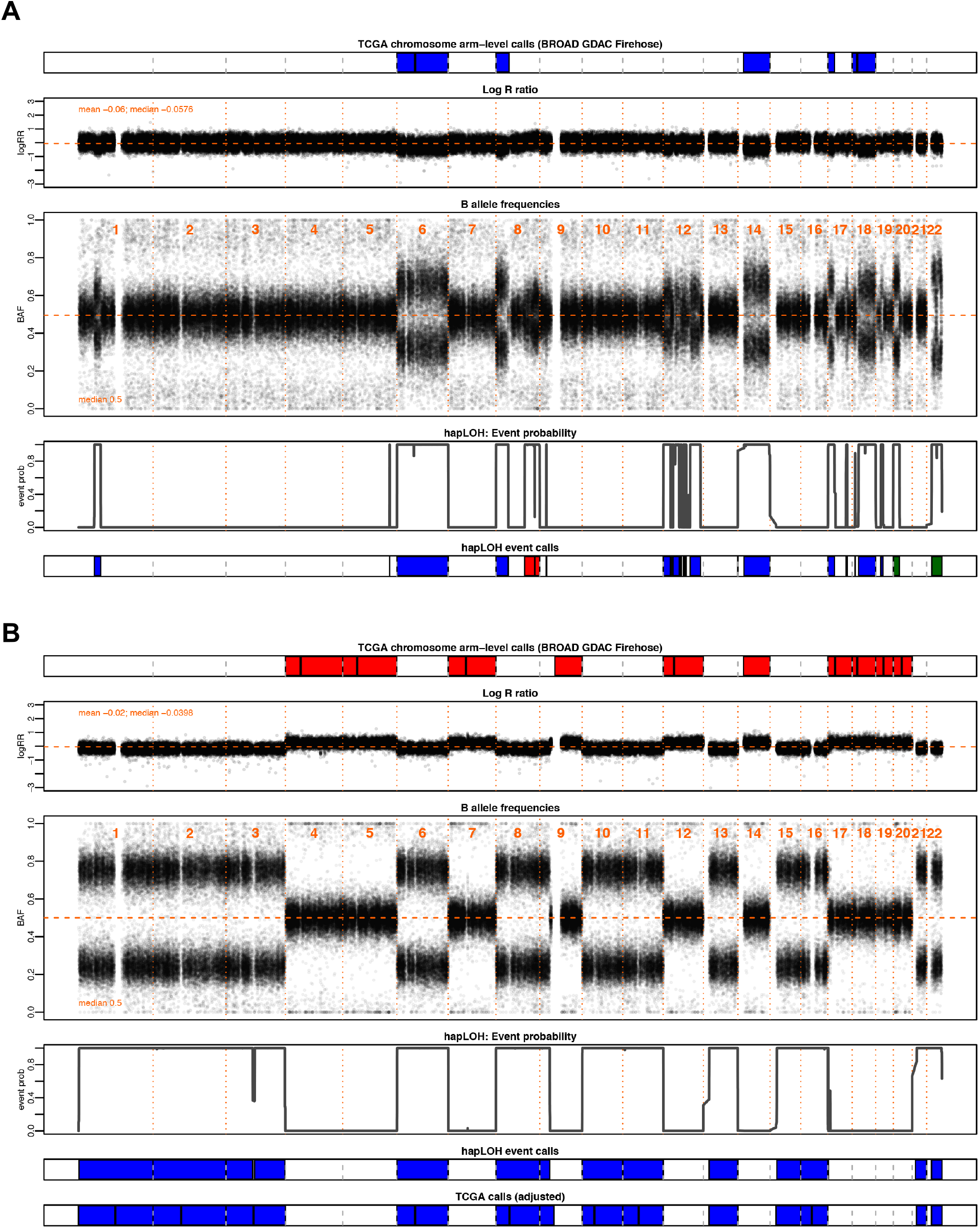
Examples highlighting the utility of a “B allele” frequency in the identification of chromosomal aberrations. Through this investigation, we aim to supplement the SCNAs in TCGA with allelic imbalance (AI) inferred from “B allele” frequency (BAF) patterns, a complementary data element to the log R ratio (LRR), from which existing calls are made, to identify additional chromosomal aberrations. Shown here are two motivating examples from pancreatic adenocarcinoma (PAAD). The tumor samples are annotated with chromosomal arm-level events downloaded from BROAD GDAC Firehose along with the BAF and LRR values at markers profiled across the genome for that individual. Below these panels are the event probabilities inferred from hapLOH using the BAF patterns, as well as classified event calls from hapLOH using a threshold-based approach from BAF and LRR deviations, for the identified event boundaries (Methods). Although all hapLOH events are shown, only chromosomal-arm level events were used for the comparison to SCNAs identified in TCGA. *(A) A PAAD tumor exhibiting overall concordance between the two call sets, with additional cnLOH events such as those on chromosome 22 and arm 20p, identified by hapLOH.* In such cases, our approach of a BAF-derived AI estimator supplements the database with additional, potentially impactful, chromosomal aberrations. *(B) A PAAD sample with discordant calls between the two approaches.* The incorporation of deviations in BAF suggest a misestimation of the normal region. The SCNAs reported in TCGA do not align with BAF deviations. However, hapLOH identifies regions based on BAF-derived AI. We apply an automated adjustment approach (Methods) for such discordant cases, the result of which is shown in the bottom-most panel. The SCNAs, after adjustment, align with deviations in BAF and thereby are concordant with hapLOH event calls. Through this approach, we address differences between the call sets and suggest methods to adjust specific cases with potentially problematic calls to aid the database with more accurate SCNAs.

In this study, we analyzed 11,074 tumor-normal pairs across 33 tumor sites in the TCGA cohort to utilize BAF as well as LRR metrics from SNP genotyping arrays to identify genome-wide patterns of allelic imbalance. Using a more sensitive approach, we were able to supplement the cohort with additional subtle chromosome-arm level SCNAs, resolve potentially conflicting cases based on prior results, and characterize the previously unknown pan-cancer landscape of chromosomal cnLOH events.

## RESULTS

Tumor genomes often exhibit high genomic instability, rendering automated identification of copy number changes challenging due to limited normal regions in the genomes which serve as a baseline for comparison. Here, we applied a sensitive haplotype-based technique to identify the landscape of chromosomal copy number changes (e.g.; gain, loss) as well as previously-uncharacterized chromosomal cnLOH events through a survey of paired tumor-normal specimens from 11,074 cases across 33 tumor types in TCGA (Table 1).

**Table 1.** Summary of allelic imbalance profiles identified across 33 tumor sites in TCGA

| Tumor site | TCGA abbreviation | Number of samples | Number of samples with AI | Samples with AI (%) | Genomic burden (Median) |
| --- | --- | --- | --- | --- | --- |
| Adrenocortical carcinoma | ACC | 90 | 90 | 100.00 | 0.6326 |
| Bladder Urothelial Carcinoma | BLCA | 411 | 408 | 99.27 | 0.5300 |
| Breast invasive carcinoma | BRCA | 1101 | 1093 | 99.27 | 0.4366 |
| Cervical squamous cell carcinoma and endocervical adenocarcinoma | CESC | 303 | 301 | 99.34 | 0.3444 |
| Cholangiocarcinoma | CHOL | 36 | 34 | 94.44 | 0.3842 |
| Colon adenocarcinoma | COAD | 462 | 455 | 98.48 | 0.3656 |
| Lymphoid Neoplasm Diffuse Large B-cell Lymphoma | DLBC | 48 | 47 | 97.92 | 0.2011 |
| Esophageal carcinoma | ESCA | 186 | 184 | 98.92 | 0.6446 |
| Glioblastoma multiforme | GBM | 545 | 539 | 98.90 | 0.2088 |
| Head and Neck squamous cell carcinoma | HNSC | 527 | 526 | 99.81 | 0.4324 |
| Kidney Chromophobe | KICH | 66 | 64 | 96.97 | 0.4115 |
| Kidney renal clear cell carcinoma | KIRC | 529 | 521 | 98.49 | 0.1731 |
| Kidney renal papillary cell carcinoma | KIRP | 288 | 281 | 97.57 | 0.1996 |
| Acute Myeloid Leukemia | LAML | 200 | 106 | 53.00 | 0.0014 |
| Brain Lower Grade Glioma | LGG | 527 | 517 | 98.10 | 0.1317 |
| Liver hepatocellular carcinoma | LIHC | 377 | 369 | 97.88 | 0.3194 |
| Lung adenocarcinoma | LUAD | 517 | 514 | 99.42 | 0.5539 |
| Lung squamous cell carcinoma | LUSC | 502 | 495 | 98.61 | 0.5921 |
| Mesothelioma | MESO | 87 | 85 | 97.70 | 0.2800 |
| Ovarian serous cystadenocarcinoma | OV | 595 | 592 | 99.50 | 0.6202 |
| Pancreatic adenocarcinoma | PAAD | 185 | 170 | 91.89 | 0.2627 |
| Pheochromocytoma and Paraganglioma | PCPG | 183 | 179 | 97.81 | 0.1543 |
| Prostate adenocarcinoma | PRAD | 496 | 465 | 93.75 | 0.0910 |
| Rectum adenocarcinoma | READ | 168 | 167 | 99.40 | 0.5100 |
| Sarcoma | SARC | 261 | 253 | 96.93 | 0.4069 |
| Skin Cutaneous Melanoma | SKCM | 472 | 469 | 99.36 | 0.4263 |
| Stomach adenocarcinoma | STAD | 442 | 436 | 98.64 | 0.4303 |
| Testicular Germ Cell Tumors | TGCT | 156 | 156 | 100.00 | 0.6504 |
| Thyroid carcinoma | THCA | 509 | 195 | 38.31 | 0.0000 |
| Thymoma | THYM | 124 | 85 | 68.55 | 0.0263 |
| Uterine Corpus Endometrial Carcinoma | UCEC | 546 | 481 | 88.10 | 0.1174 |
| Uterine Carcinosarcoma | UCS | 55 | 55 | 100.00 | 0.5941 |
| Uveal Melanoma | UVM | 80 | 79 | 98.75 | 0.1628 |

### Pan-cancer allelic imbalance burden

Our method identified at least one AI event in 10,411 cases (94%), with a median genomic AI burden (defined as the percentage of a sample’s genome exhibiting AI) of 32%. We further assessed the patterns of genomic AI burden for each tumor site independently, to account for the site-specific molecular complexities and variation in number of samples across tumor types, displayed in Figure 2 and Table 1. Testicular germ cell tumors (TGCT), esophageal carcinoma (ESCA), adrenocortical carcinoma (ACC) and ovarian carcinoma (OV) exhibited high overall genomic AI burdens, with medians for each exceeding 60%. Uterine carcinosarcoma (UCS), lung squamous cell carcinoma (LUSC), and lung adenocarcinomas (LUAD) also exhibited high genomic AI burden, with medians of over 50%. At low end of the burden spectrum were thyroid carcinoma (THCA), acute myeloid leukemia (LAML), thymoma (THYM), prostate adenocarcinoma (PRAD), all of which exhibited a median genomic AI burden below 10%. Each event type also showed varied patterns for genomic burdens across tumor types. TGCT, UCS, ESCA and OV showed highest median genomic burdens for gains, while ACC, KICH, OV, LUSC and UCS showed high rates of genomic burdens for losses. The different tumor sites also exhibited different patterns of enrichment of the three event types. While some cancers, such as KICH and ACC, showed pronounced and preferential enrichment of loss events, tumors such as KIRP and TGCT showed high gain burdens (Supplementary Figure 1). The relative abundance of cnLOH was overall lesser than gains and losses, and spanned smaller proportions of the genome; the highest rates of cnLOH genomic burdens were observed in TGCT, ESCA, LUSC, UCS and OV.

**Figure 2.**
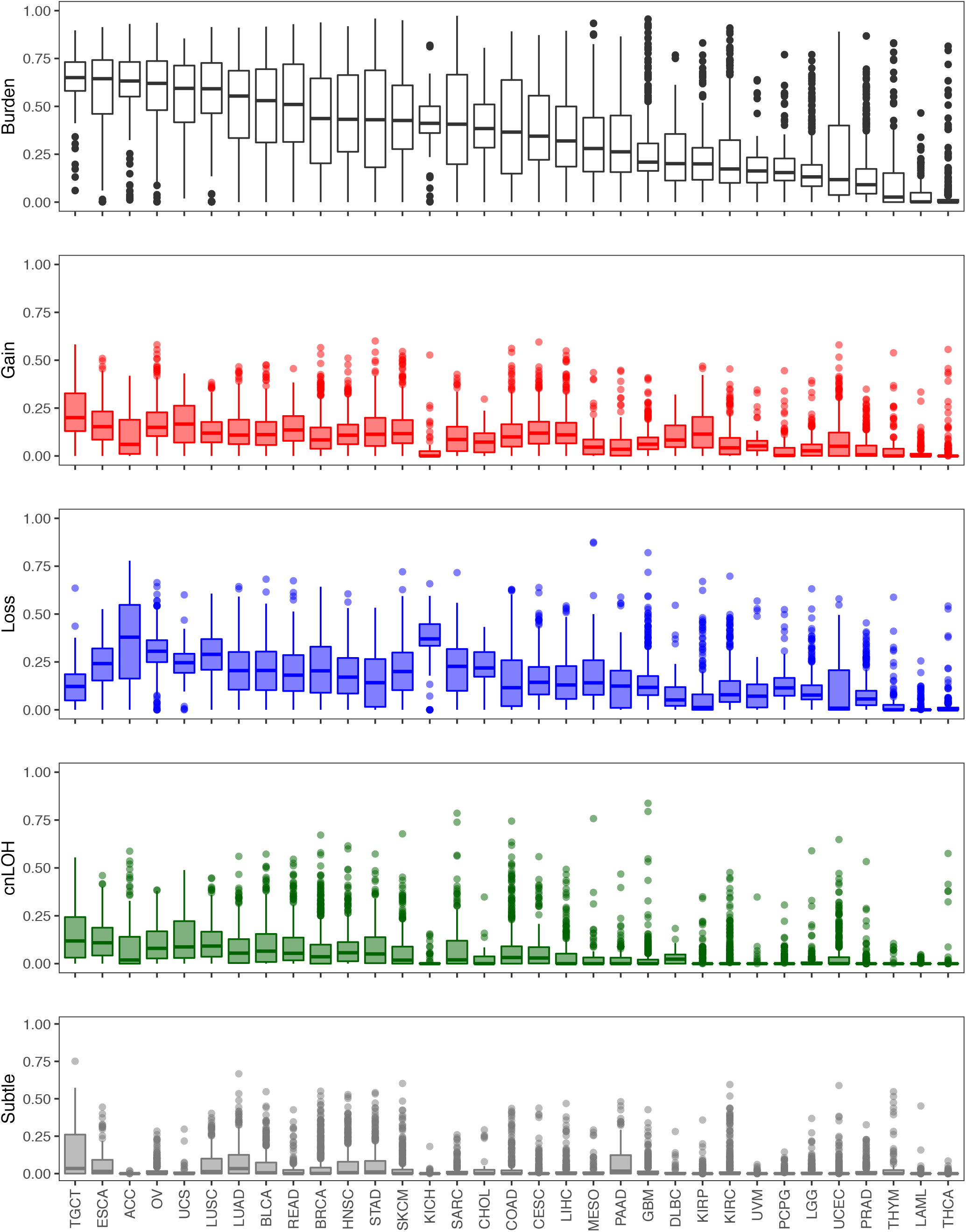
The distribution of genomic allelic imbalance burden across 33 tumor sites in TCGA. We identified regions of allelic imbalance (AI) across 11,074 tumor samples across 33 tumor sites in TCGA. Boxplots representing the distribution of overall genomic AI burden, defined as a percent of the genome exhibiting evidence of AI, are shown for each tumor site. The distribution of genomic burdens for each event type are also shown (red: gain, blue: loss, green: cnLOH, grey: subtle, unclassifiable events). The tumors are ordered by their median overall genomic burden.

### Landscape of chromosome arm-level copy number changes across tumor sites

Since our method is tuned for the detection of mega-base scale chromosomal changes such as those that span an entire chromosome or chromosome arm, we examined in greater depth chromosomal arm events across the 33 tumor types (Figure 3). At the arm level, our method identified 121,645 events in 10,004 tumor samples, of which 32,925 were gains and 57,161 were losses (Supplementary Table 1). Among these, the most common pan-cancer chromosome arm event occurred on 17p (Figure 3). Although 17p events were common among multiple tumor sites including ACC, KICH, COAD, LUAD, LUSC, PAAD, ESCA and BRCA, some tumor sites did not show an enrichment for allelic imbalance on 17p, such as GBM, KIRC, THCA, UVM an PRAD (Figure 3). Among cases that showed copy number changes on 17p, most comprised loss events; KIRP was the only tumor site that showed an abundance of 17p gains. Events on 8p and 3p were also prevalent across multiple tumor sites (Figure 3). While most cancer types showed a loss of 8p, LAML and UVM exhibited predominantly gain events; STAD, UCEC, and COAD showed mixed event types on 8p (Figure 3). Particularly in PRAD, 8p loss events were predominant with rest of genome being relatively stable, showing limited events in the rest of the genome such as 8q gain and 18q loss. As with 17p events, loss of 3p occurred across many tumor sites (Figure 3). Particularly in KIRC, 3p loss events seemed to be a driver, with rest of the genome showing very limited evidence for chromosomal instability (Figure 3). Loss of 3p was also prevalent in UVM, LUAD, LUSC, HNSC, CHOL and CESC (Figure 3). Amplification of 8q was the most frequent pan-cancer gain event, showing high occurrence in multiple tumor sites including UVM, LAML, COAD, HNSC, STAD, UCEC, SKCM and LIHC (Figure 3). The second most prevalent amplification was identified on chromosome 7p, particularly in KIRP, DLBC, COAD, GBM, SKCM and STAD.

**Figure 3.**
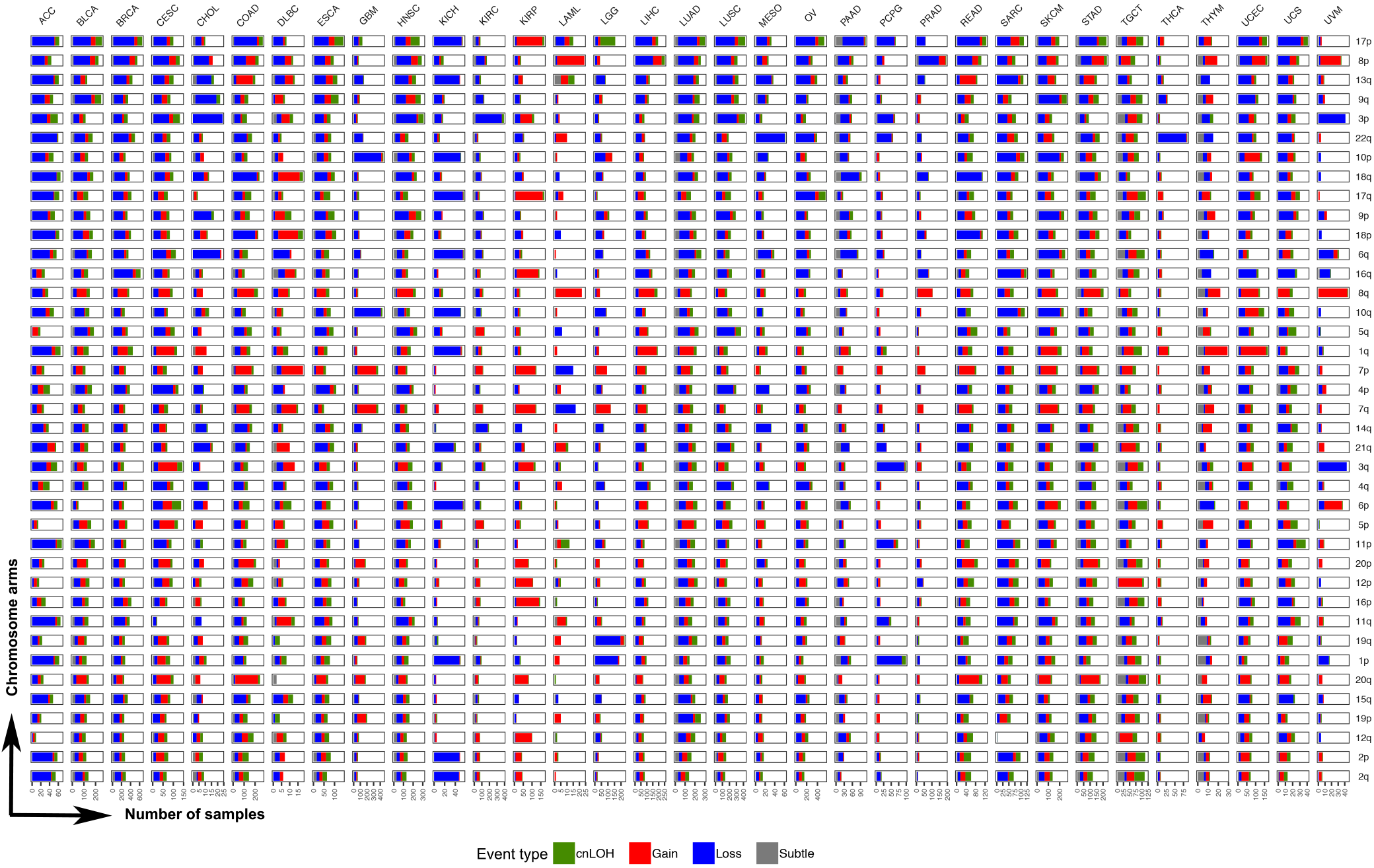
Pan-cancer patterns of chromosomal arm-level allelic imbalance. We identified allelic imbalance events that spanned at least 70% of the genome as chromosome-arm level events (Methods). The distribution of these events across all non-acrocentric autosomal chromosome arms (n=39) are shown for the 33 tumor sites studied. The chromosome arms are sorted by their overall frequency of event occurrence across all tumor sites. For each tumor site, a stacked barplot of the number of tumor samples with event type-specific chromosome-arm level events are shown for all 39 chromosome arms investigated.

Amplification of 1q was also observed across many tumor sites; LUAD, LIHC, CESC, UCEC, SKCM and THYM showed relatively high occurrences of 1q gain (Figure 3). Similarly, an amplification of 20q seemed to be prevalent in gastrointestinal tumors COAD, READ and STAD. (Figure 3). In contrast, chromosome arms 2p and 2q were the least altered across tumor types; a predominant loss of 2p and 2q was observed only in ACC and KICH datasets, both of which consisted of very few cases.

Tumor sites could broadly be classified into two categories, based on the distribution of chromosomal arm events across the genome. Tumor sites such as UVM, THCA, KIRC, LGG, LAML and PRAD showed a significant enrichment of at most a single or very few chromosome arm events with the remaining parts of the genome being stable (Figure 3). For example, three tumor sites showed single arm events that dominated the AI profile in those tumors such as 22q loss in THCA, 3p loss in KIRC and 8p loss in PRAD tumors. LGG tumors showed a significant enrichment of 1p and 19q loss events, consistent with the known phenotype of 1p/19q codeletion in LGGs. In contrast to LGGs, the other class of brain tumors, GBMs, exhibited a different allelic imbalance profile. In GBM tumors, the frequent events included the loss of chromosome 10 and gain of chromosome 7 (Figure 3). LAML tumors, although from a small dataset, showed high prevalence of chromosome 8 gains and to a lesser extent chromosome 7 loss events (Figure 3). UVM tumors also seemed to exhibit relatively few chromosomal arm events that spanned losses of chromosome 3 and 6q, and gains of chromosome 8 gains and 6p. PCPG also showed few chromosome arms under allelic imbalance, the most frequent being loss events on 1p and 3q. In contrast to these tumor sites that showed lower overall allelic imbalance burden, which was often accompanied by the enrichment of a single or few chromosome arm events, tumor sites such as LUAD, LUSC, BRCA, ESCA, ACC, TGCT, STAD, SKCM and SARC showed genome-wide allelic imbalance patterns involving multiple chromosome arms. Tumor sites could also be classified based on the enrichment of a specific event type among the allelic imbalance events detected. For example, KIRP tumors seemed to be primarily driven by gain events across the genome. In contrast, tumor sites such as KICH, PCPG, STAD and PAAD showed genome-wide enrichment of losses (Figure 3, Supplementary Figure 1). These results aid in understanding the role of different chromosomal changes and burdens in driving the development of different tumor types, based on their sites of origin.

### Copy neutral loss of heterozygosity patterns across tumor sites

We analyzed TCGA tumor samples to identify deviations in the expected 1:1 allelic ratios at germline heterozygous loci, a by-product of which is accurate detection of regions of copy neutral loss of heterozygosity (cnLOH) that result in allelic ratios of 2:0 or 0:2 in the cells with the chromosomal mutation. Standard methods that rely on identifying copy number changes from LRR (total allele) intensities will miss this particular class of chromosomal aberrations. Our allelic imbalance annotation of the 33 tumor sites in TCGA that includes cnLOH status, which, to date, has not been comprehensively characterized. Our methods identified 20,454 cnLOH arm-level events across 5,222 cases in TCGA (Supplementary Table 1). Figure 4 shows the distribution of chromosome arm-level cnLOH across the genome for each tumor site. Among these, TGCT and ESCA showed the highest rates of cnLOH (by burden and arms; Figure 1, Figure 4). Chromosome 17 showed the highest rates of cnLOH across tumor sites (Figure 4). Chromosome arm 3p, 6p and 2p also showed high rates of cnLOH events across tumor sites (Figure 4).

**Figure 4.**
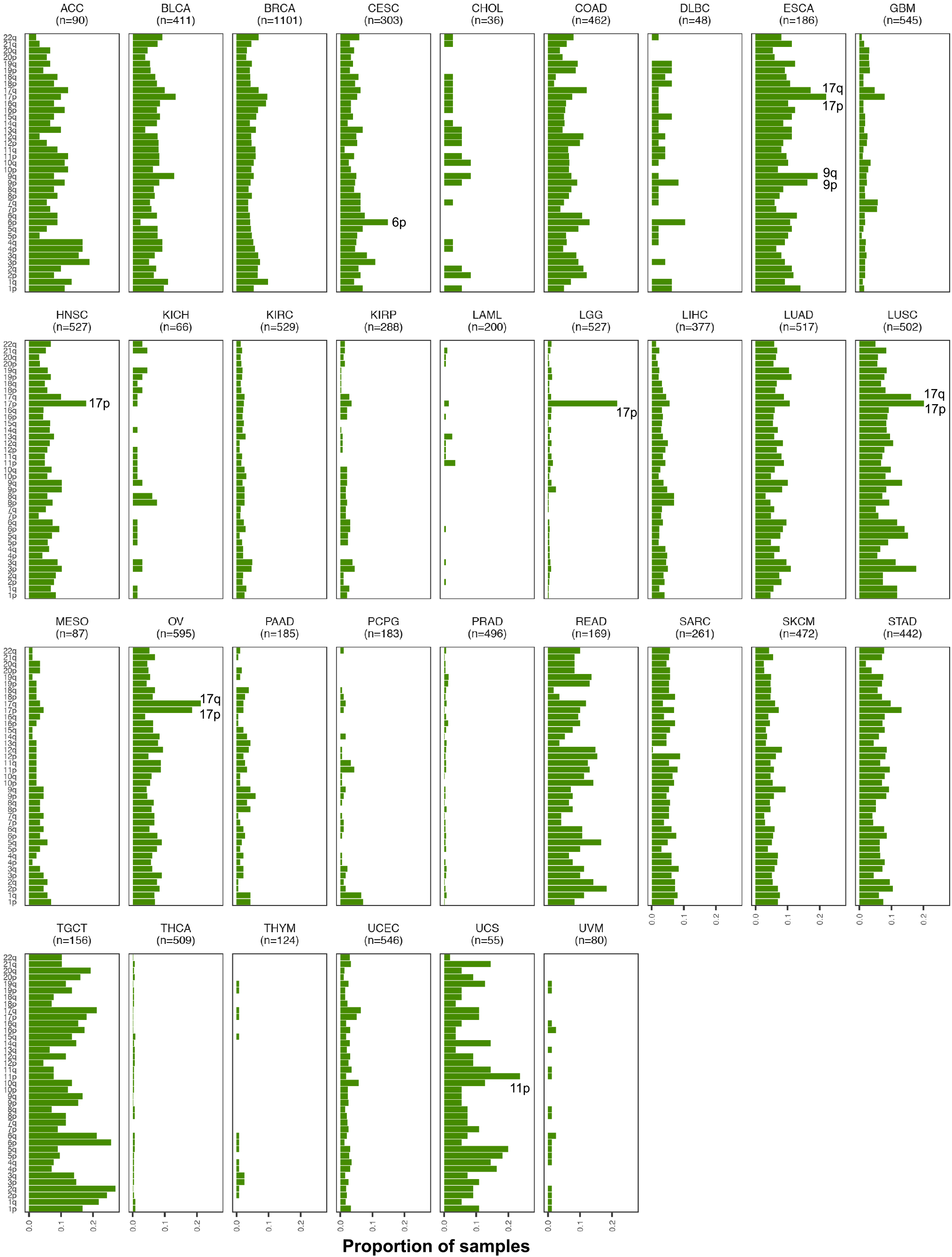
Landscape of chromosomal arm-level copy-neutral loss of heterozygosity across TCGA. The allelic imbalance-based approach implemented here allowed for the identification of the previously unexplored landscape of copy-neutral loss of heterozygosity (cnLOH) patterns across all tumor sites in TCGA. Shown here are bar plots representing the proportion of samples exhibiting cnLOH events across chromosome arms in the genome, for each tumor site. Enriched pan-cancer cnLOH events as well as site-specific cnLOH events are highlighted within the respective tumor site plots.

We interrogated in greater depth the chromosome arm-level cnLOH that showed enrichment within specific tumor sites (Figure 4). For example, LGG tumors showed a pronounced presence of cnLOH on 17p (21.6% of tumors, Figure 4), an observation that would be completely missed in copy number detection approaches that use LRR intensities only. In addition, cases that exhibited 17p cnLOH in LGGs (n=114) were found to be mutually exclusive of cases that exhibited the 1p/19q co-deletion (n=160). Copy neutral LOH of 17p were also prevalent in ESCA (22%), LUSC (20.1%), OV (18.5%) and HNSC tumors (17.8%) (Figure 4). These tumor sites also showed evidence for cnLOH events on the q arm of chromosome 17 (*BRCA1, NF1*). 6p (*DAXX*) also showed high rates of cnLOH, particularly in CESC tumors (14.8% of tumors). In UCS, amidst high rates of cnLOH across the genome, 11p (*WT1*) cnLOH events were pronounced (23.6% of tumors). We also observed tumor sites with multiple cnLOH events across the genomes, i.e., without having any visually observable specificity for particular chromosome arm events. In addition to TGCT and ESCA, mentioned above, READ, LUAD, SKCM and STAD exhibited cnLOH across the genome. These results suggest the importance of investigating cnLOH events to better understand their role in oncogenesis and determine the molecular mechanisms leading to these chromosomal aberrations across all tumor sites.

### Recalibration of TCGA copy number profiles

Our allelic imbalance analysis provides a set of potential SCNAs based on data orthogonal to existing LRR-based SCNA calls. We sought to interrogate these sets of calls for consistency to potentially aid automated algorithms for SCNA identification. We assessed concordance in allelic imbalance and TCGA-identified SCNAs for 10,680 tumor samples, among which a Pearson correlation could be calculated for 9,364 samples (Supplementary Table 2). Overall, 7,577 showed showed putative consistency (positive correlation) between the two sets (Supplementary Table 2). For 1,787 cases, a surprising negative correlation existed between the calls. A closer inspection of these revealed a strong negative correlation (Pearson correlation, −0.7417) between the overall genomic burden of allelic imbalance and the concordance of the two call sets, i.e., samples that exhibited high overall allelic imbalance burdens tended to show patterns of discordance between the two calls sets. This trend was consistent across all tumor sites (Supplementary Figure 2).

We identified a subset of 1653 cases that exhibited a negative correlation between the two call sets as well as a high overall genomic AI burden (>=50% of the genome). These cases are listed in Supplementary Table 3. Figure 5 displays the proportion of these discordant samples across tumor sites. The highest proportion of negatively correlated samples were observed in TGCT (64.7%) and ACC (54.4%) consisting of a total of 150 and 90 profiled cases respectively. UCS also showed a high proportion of discordant calls (37%) from a total of 50 cases profiled. All three datasets were relatively small, which might be inflecting the percentage of discordant samples. However, we also observed relatively high levels of discordance, greater than 25% of the samples, in larger studies such as ESCA (30.4%), READ (27.8%), OV (27.6%), BLCA (26.2%) and LUAD and LUSC (24%) (Supplementary Table 2). In contrast, we did not identify any discordant calls in LAML (n=191). Other tumor sites that exhibited very low percent of discordant call sets were THCA (1.2%), LGG (1.2%), PRAD (1.2%) and DLBC (2.1%) (Supplementary Table 2).

**Figure 5.**
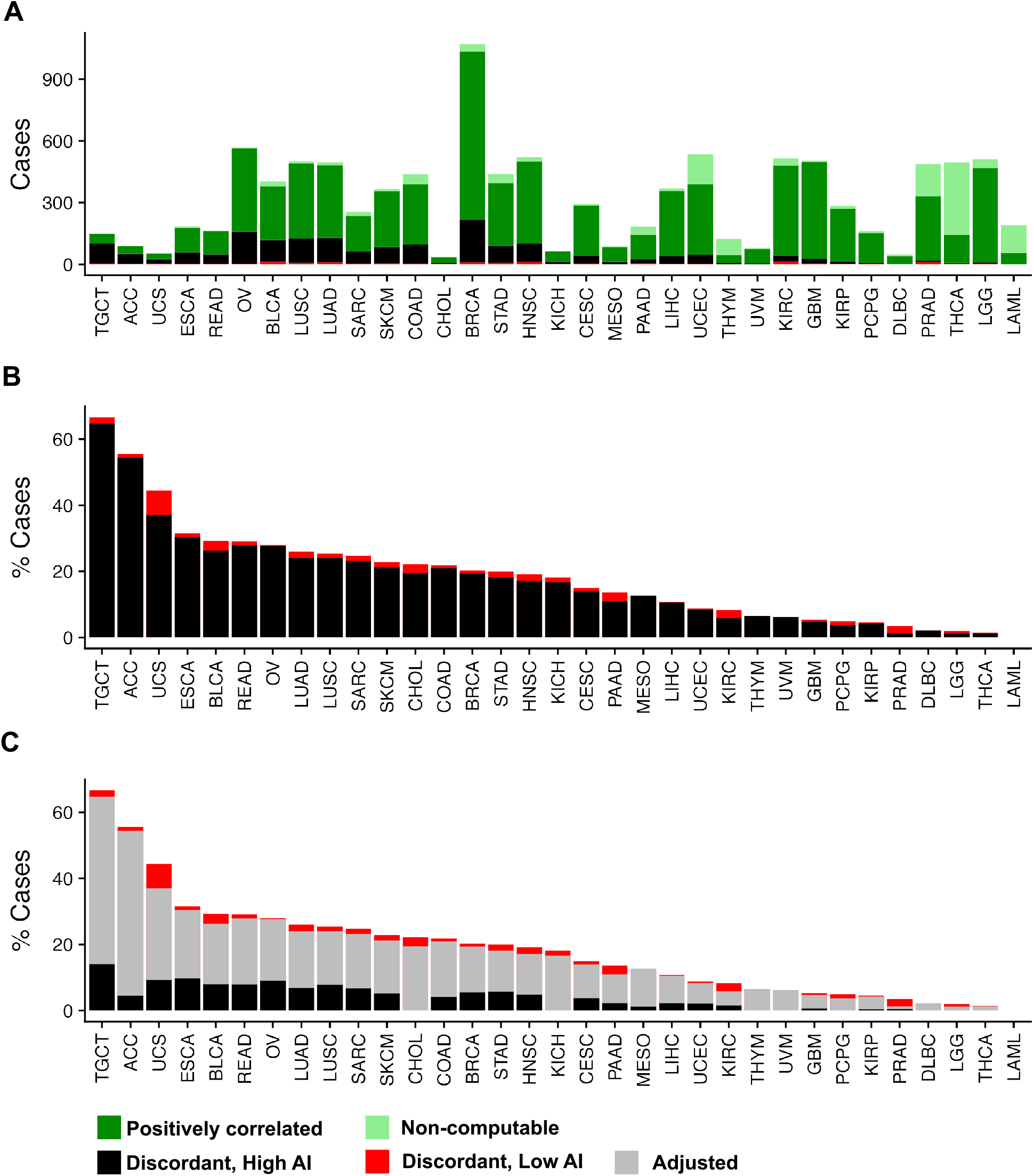
Identification and adjustment of putative discordant samples between SCNAs inferred from hapLOH with those reported in TCGA. A total of 10,680 tumor samples were assessed for concordance between the two call sets (Methods). (A) The distribution of positively correlated (dark green) and negatively correlated (black and red) samples for each tumor site are shown as a stacked bar plot. The negatively correlated samples are further divided into two categories, based on their overall allelic imbalance (AI) genomic burden, with cases showing at least 50% of their genome aberrant termed as high AI (black), with the remaining cases annotated as low AI (red). Samples for which correlation was non-computable, based on the absence of at least one event in each call set are also shown (light green). These cases comprise a vast majority of samples that exhibited no evidence of SCNAs in either of the call sets. (B) A stacked bar plot of the distribution of negatively correlated samples (high AI and low AI), as a percentage of the total number of samples profiled for each tumor site is shown. (C) An automated adjusted procedure was applied on cases identified to be ‘negatively correlated, high AI’ (black). A stacked bar plot of the distribution of samples after applying the adjustment procedure are shown across all tumor sites, with the percent of adjusted cases shown in grey. Tumor sites in all three panels are ordered by their overall percent of putative discordant samples.

Within this set of samples for which our allelic imbalance calls were discordant from TCGA SCNAs, we suspected a potential cause was that the level of aneuploidy was sufficiently high such that it was difficult to auto-detect the baseline LRR in identifying the SCNAs. Under the assumption that normal copy number regions will not typically exhibit allelic imbalance, we further sought to bolster the TCGA SCNA calls in this discordant set by re-establishing the baseline LRR regions and recalibrating the SCNA values. After adjusting the calls, a positive concordance was achieved in 1224 (of 1653) samples. The performance of our automated adjustment protocol appeared to vary by tumor site (Figure 5, Supplementary Table 2). For example, in tumor sites such as LGG, THCA, PCPG and THYM, our approach successfully adjusted all discordant cases to achieve a positive correlation (Supplementary Table 2). ACC and MESO also achieved high rates of adjustment of over 90% of the potentially problematic cases (Supplementary Table 2). Across tumor sites, after performing the automated adjustment, the percent of negatively correlated samples that remained was drastically lower. For example, the percent of negatively correlated samples reduced from 54.4% to 4.4% in ACC, 23.1% to 6.6% in SARC, 21.2% to 5.1% in SKCM, 20.9% to 4.1% in COAD. However, even after adjustment, some tumor sites showed an appreciable number of discordant samples. For example, TGCT showed negative correlation in 14% cases after adjustment; similarly ESCA, UCS and OV also showed rates of 9.8%, 9.2% and 8.9% respectively, after adjustment (Supplementary Table 2). Nonetheless, our approach was able to significantly reduce the rates compared to the trends observed before adjustment across all tumor sites.

## DISCUSSION

Acquired chromosomal aberrations such as deletions, duplications and copy-neutral loss-of-heterozygosity (cnLOH) serve as hallmarks of tumorigenesis. Large (megabase-scale) alterations span multiple heterozygous markers and thus result in deviations from the expected one-to-one allelic ratio, thereby leading to allelic imbalance (AI). The SCNA pipeline of the TCGA consortium reports genomic regions and their segment mean copy number estimates from SNP genotyping arrays. However, detection and accurate identification of SCNAs in this resource are limited by the strict reliance on summaries of total intensities. Here we sought to address these limitations by modeling the allele-specific intensities (B allele frequencies; BAFs), leveraging an expectation that the perturbations in the BAFs within most SCNAs will occur in a manner consistent with the allelic patterns on the inherited chromosomes, ie. the haplotypes. Our haplotype-based approach, by jointly modeling the BAFs, more powerfully detects SCNAs that lead to patterns of AI as well as AI induced by cnLOH. In this manuscript we provided a summary of these AI findings in 11,074 samples from all available tumor sites.

A recent study summarized pan-cancer chromosomal aberrations and correlated chromosomal arm aneuploidy to somatic point mutations and expression of immune signaling genes (Taylor et al. 2018). Chromosomal copy number events identified in our present study corroborated with their findings, such as the high prevalence of loss events on 17p and 8p, as well as gains of 8q across tumor sites. Similarly, both studies identified chromosome 2 as the least aberrant across tumor sites. Our allelic imbalance burden derived estimates were also similar to the aneuploidy scores across TCGA types (Taylor et al. 2018); however we report higher burdens in ESCA and OV. Another pan-cancer atlas of the TCGA tumor sites identified specific clusters of tumors based on the extent of aneuploidy and the type of events (Hoadley et al. 2018). We identified similar patterns from our allelic imbalance derived chromosome arm-level aberrations. Both studies identified a subset of low burden sites that included PRAD, THYM, LAML and THCA. Our results also identified the enrichment of 13q gain and chromosome 18 loss in gastrointestinal tumors (COAD, READ, and STAD); in addition to these events, we also observed high rates of gains on chromosome 20 in these tumors. Similarly, we also identified the enrichment of chromosome 7 gains and chromosome 10 losses in GBM tumors. In comparison to both these previous studies (Taylor et al. 2018; Hoadley et al. 2018), we also provide a complementary landscape of pan-cancer cnLOH events that opens a window of opportunities to investigate the importance of these cnLOH events in tumorigenesis. A striking result observed in our study was the recurrent cnLOH of 17p across multiple tumor sites. Future studies investigating the differences between previously reported pan-cancer 17p deletions and 17p cnLOH patterns reported here, as well as their roles with respect to the presence of *TP53* variants or mutations can aid in better understanding the dynamics of tumors that exhibit multi-hit aberrations. Based on recent evidence (Liu et al. 2016), it is also plausible that these pan-cancer 17p loss and cnLOH affect a combination of genes, extending beyond the effects on the *TP53* tumor suppressor gene.

Our study also showcased high rates of previously unknown chromosomal aberrations across multiple tumor sites. For example, in KIRP tumors that were predominantly driven by gain events, our findings aligned with previously identified high frequency gains on chromosome 7 and 17, but we additionally identified a higher proportion of chromosome 16 gains than previously reported (Cancer Genome Atlas Research Network et al. 2016). Similarly, in UCS tumors, we identified recurrent loss and cnLOH events of 17p as well as novel recurrent cnLOH of 11p. Given the prevalence of TP53 mutations in UCS (Cherniack et al. 2017), cnLOH of 17p may be part of somatic two-hit mechanisms of mutagenesis in UCS tumors. In HNSC tumors, the highest rates of chromosomal changes were previously reported on 8p and 3p (Cancer Genome Atlas Network 2015). Our method not only identified these events but also detected high rates of 17p loss and cnLOH events in these tumors. In this way, findings from our study supplement the current database of chromosome-arm SCNAs across multiple tumor sites, through the detection of additional chromosomal aberrations.

It is noteworthy that cnLOH events we identified to be enriched in our present investigation across multiple tumor sites have been shown to be prognostically relevant in independent studies. For example, we observed recurrent 17p cnLOH in LGGs in addition to the well-known 1p/19q codeletion; cnLOH of 17p, as well as its mutual exclusivity with 1p/19q codeletion, has been previously shown to be a potential marker in independent cohorts of gliomas (Labussière et al. 2016; Idbaih et al. 2012; Suzuki et al. 2015). Similarly, our finding of 6p cnLOH in CESC tumors corroborates previous studies that have identified LOH on 6p21.2 that correlated with recurrence of cervical carcinoma after radiotherapy (Harima et al. 2000). This suggests that cnLOH events presented in our study have the potential to serve as prognostic markers for the detection or prediction of recurrence across tumor sites. Although unlikely to act as a driver mutation alone, with the possible exception of differential germline configurations, the high rates of cnLOH presented here highlight the importance of the acquired mutation on the preserved chromosome in a region of cnLOH. Thus, regions of cnLOH warrant additional study to better assess the multiple steps in tumor pathogenesis.

In addition to augmenting the TCGA with allelic imbalance annotation, which helps to address previously undetected SCNAs and cnLOH, our results also allow for improving the accuracy of some of the existing SCNA calls. Tumor samples that displayed conflicting TCGA SCNA and allelic imbalance profiles suggested incorrectly-estimated “normal regions” for obtaining baseline LRR signal intensities. These observations, with the corresponding wide user base of the TCGA repository, motivated us to develop an automated procedure to identify and adjust these putative problematic cases. Through our efforts in this study, we list these samples as well as provide a simple adjustment procedure to renormalize the data for potentially improved SCNA detection in a select group of samples.

Although our methods to identify SCNAs and cnLOH, as well as adjust previously-identified SCNAs, provides or augments a useful resource for pan-cancer genomic investigations, they do not come without limitations. First, our approach relies on deviations in allelic ratios at germline heterozygous sites to identify regions of allelic imbalance. As such, we do not attempt here to identify *balanced* SCNAs (e.g. both copies of a chromosome lost or gained in a proportion of cells). Second, in line with the requirement of altered allelic ratios, our approach is at least slightly better at detecting loss or cnLOH events that results in a more severe change in allelic ratios (i.e. 1:0 or 2:0), in comparison to gains (e.g., one copy gains that results in a ratio of 1:2). Third, although our algorithm provides sensitive identification of regions of allelic imbalance, we currently implement a naive thresholding-based classification to annotate events as loss, gain and cnLOH. This results in detection of subtle events that robustly show a signal for allelic imbalance but are unclassifiable using our current LRR threshold-based approach. Nonetheless, these additional events supplement the current repository of copy number alterations, and future computational methods to jointly utilize BAF and LRR in the statistical estimation of allelic imbalance might overcome this issue. Lastly, our prototype methodology to systematically adjust mean segment copy numbers will in some instances suffer from over-corrections, as the procedure is applied genome-wide without appropriate tuning. Conversely, the approach may under-correct in situations where regions identified as normal by hapLOH contain both gains and losses as identified by TCGA; in such cases, our approach will not identify a deviant copy number from which to correct. However, these limitations notwithstanding, our work highlights the importance of integrating multiple data types (“B allele” frequency and log R ratio) for more robust automated inference procedures.

While we acknowledge further enhancements to what we have presented here may be possible with greater statistical sophistication, results presented here have the potential to support current methods and improve downstream analyses, including clinical evaluations and identification of complex SCNA-derived signatures. We hope that enhanced annotation of this widely-used and valuable public resource may support new hypotheses of chromosomal instability in tumorigenesis.

## METHODS

### Dataset

The Level 1 raw CEL files from Affymetrix Genome-Wide Human SNP Array 6.0 profiling of 11,074 paired tumor-normal samples across 33 cancer sites in The Cancer Genome Atlas (TCGA) were downloaded from the Genomic Data Commons (GDC) data portal, along with available clinical annotations. The cohort comprised a majority of primary tumors (n=10,680), with a few metastatic specimens (n = 394; SKCM accounting for 368 of these samples). SNP metrics including genotypes, “B allele” frequency (BAF) and log R ratio (LRR) were obtained from the Birdsuite software (Korn et al. 2008).

### Pan-cancer allelic imbalance profiles using hapLOH

For each tumor sample, the corresponding control sample (blood or tumor adjacent normal tissue) was paired and statistical reconstruction of haplotypes was performed using MaCH (Li et al. 2010). The human genome build hg19 (GRCh37) was used as the reference. The phased genotypes as well as the BAFs were supplied as inputs to run hapLOH using default parameters. The resulting regions of allelic imbalance (AI) as inferred from hapLOH were then characterized based on the extent of BAF and LRR deviations for each event region. Regions with LRR deviation greater than or equal to 0.05 were classified as gains while those with LRR deviations below −0.05 were classified as losses. Of note, AI events with LRR>=0.08 and less than 2Mb in length were excluded as likely inherited duplications. The remaining events were characterized as copy-neutral loss of heterozygosity if median BAF deviations were >0.1. The events that we were unable to be characterized into these different types were deemed to be too subtle, due to low mutant cell fraction, for annotation and labeled as “Subtle”. The allelic imbalance events spanning more than 70% of the genome were considered chromosomal-arm level events, while the remaining were retained as focal events. For each tumor sample, the percent of its genome under allelic imbalance was used as a measure of its genomic AI burden.

### TCGA pan-cancer copy number profiles

The processed chromosome arm-level copy number events files were downloaded from Broad GDAC Firehose. The latest analyses version at the time of download was dated *2016_01_28*. The *broad_values_by_arm.txt* files under each TCGA study were then processed into *bed* formatted files using the specified amplification/deletion threshold of 0.1. These results were compared with the allelic imbalance derived copy number events identified by hapLOH for the same individuals.

### Identification of putative problematic calls in TCGA

Allelic imbalance events identified in each tumor sample were reassessed at a chromosome-arm level, by concatenating events on each arm to identify specific chromosome-arms that exhibited events at a fraction 0.7 of the arm, using a custom script. For the purpose of SCNA comparisons, cnLOH events and subtle unclassified events were excluded from this analysis. For every marker genotyped in the array, the presence (or absence) of an event spanning the marker in both TCGA and hapLOH derived event calls were annotated as 1 (or 0) respectively. A Pearson correlation coefficient was computed from all markers. Samples with a negative correlation were identified as discordant and potentially problematic.

### Automated adjustment of potentially problematic calls in TCGA

For each of the negatively correlated tumor samples identified through the procedure described above, the normal region, as determined by hapLOH was identified. Events reported by TCGA within these normal regions, as well as those that were identified as normal by both methods were identified. A new weighted median copy number was calculated from these events, weighted by the length of the event. The original calls made by TCGA were recalibrated using this newly determined normal copy number. Using the same specifications as before, i.e. a amplification/deletion threshold of 0.1, the new set of chromosome-arm event calls were reclassified. A correlation between these adjusted TCGA SCNA calls and hapLOH derived SCNAs was calculated in a manner explained in the previous section.

## DATA ACCESS

Not applicable. All processed files and results have been provided as part of the Supplementary material accompanying this manuscript.

## REFERENCES

Cancer Genome Atlas Network. 2015. Comprehensive genomic characterization of head and neck squamous cell carcinomas. Nature 517: 576–582.

Cancer Genome Atlas Research Network, Linehan WM, Spellman PT, Ricketts CJ, Creighton CJ, Fei SS, Davis C, Wheeler DA, Murray BA, Schmidt L, et al. 2016. Comprehensive Molecular Characterization of Papillary Renal-Cell Carcinoma. N Engl J Med 374: 135–145.

Cherniack AD, Shen H, Walter V, Stewart C, Murray BA, Bowlby R, Hu X, Ling S, Soslow RA, Broaddus RR, et al. 2017. Integrated Molecular Characterization of Uterine Carcinosarcoma. Cancer Cell 31: 411–423.

Harima Y, Harima K, Sawada S, Tanaka Y, Arita S, Ohnishi T. 2000. Loss of heterozygosity on chromosome 6p21.2 as a potential marker for recurrence after radiotherapy of human cervical cancer. Clin Cancer Res 6: 1079–1085.

Hoadley KA, Yau C, Hinoue T, Wolf DM, Lazar AJ, Drill E, Shen R, Taylor AM, Cherniack AD, Thorsson V, et al. 2018. Cell-of-Origin Patterns Dominate the Molecular Classification of 10,000 Tumors from 33 Types of Cancer. Cell 173: 291–304.e6.

Hoadley KA, Yau C, Wolf DM, Cherniack AD, Tamborero D, Ng S, Leiserson MDM, Niu B, McLellan MD, Uzunangelov V, et al. 2014. Multiplatform analysis of 12 cancer types reveals molecular classification within and across tissues of origin. Cell 158: 929–944.

Hutchinson L. 2017. Biomarkers: Aneuploidy and immune evasion — a biomarker of response. Nat Rev Clin Oncol 14: 140.

Idbaih A, Ducray F, Dehais C, Courdy C, Carpentier C, de Bernard S, Uro-Coste E, Mokhtari K, Jouvet A, Honnorat J, et al. 2012. SNP array analysis reveals novel genomic abnormalities including copy neutral loss of heterozygosity in anaplastic oligodendrogliomas. PLoS One 7: e45950.

Knijnenburg TA, Wang L, Zimmermann MT, Chambwe N, Gao GF, Cherniack AD, Fan H, Shen H, Way GP, Greene CS, et al. 2018. Genomic and Molecular Landscape of DNA Damage Repair Deficiency across The Cancer Genome Atlas. Cell Rep 23: 239–254.e6.

Korn JM, Kuruvilla FG, McCarroll SA, Wysoker A, Nemesh J, Cawley S, Hubbell E, Veitch J, Collins PJ, Darvishi K, et al. 2008. Integrated genotype calling and association analysis of SNPs, common copy number polymorphisms and rare CNVs. Nat Genet 40: 1253–1260.

Labussière M, Rahimian A, Giry M, Boisselier B, Schmitt Y, Polivka M, Mokhtari K, Delattre J-Y, Idbaih A, Labreche K, et al. 2016. Chromosome 17p Homodisomy Is Associated With Better Outcome in 1p19q Non-Codeleted and IDH-Mutated Gliomas. Oncologist 21: 1131–1135.

Liu Y, Chen C, Xu Z, Scuoppo C, Rillahan CD, Gao J, Spitzer B, Bosbach B, Kastenhuber ER, Baslan T, et al. 2016. Deletions linked to TP53 loss drive cancer through p53-independent mechanisms. Nature 531: 471–475.

Li Y, Willer CJ, Ding J, Scheet P, Abecasis GR. 2010. MaCH: using sequence and genotype data to estimate haplotypes and unobserved genotypes. Genet Epidemiol 34: 816–834.

Lourenço N, Hélias-Rodzewicz Z, Bachet J-B, Brahimi-Adouane S, Jardin F, Tran van Nhieu J, Peschaud F, Martin E, Beauchet A, Chibon F, et al. 2014. Copy-neutral loss of heterozygosity and chromosome gains and losses are frequent in gastrointestinal stromal tumors. Mol Cancer 13: 246.

Melcher R, Hartmann E, Zopf W, Herterich S, Wilke P, Müller L, Rosler E, Kudlich T, Al-Taie O, Rosenwald A, et al. 2011. LOH and copy neutral LOH (cnLOH) act as alternative mechanism in sporadic colorectal cancers with chromosomal and microsatellite instability. Carcinogenesis 32: 636–642.

Negrini S, Gorgoulis VG, Halazonetis TD. 2010. Genomic instability — an evolving hallmark of cancer. Nat Rev Mol Cell Biol 11: 220–228.

Nichols CA, Gibson WJ, Brown MS, Kosmicki JA, Busanovich JP, Wei H, Urbanski LM, Curimjee N, Berger AC, Gao GF, et al. Loss of heterozygosity of essential genes represents a widespread class of potential cancer vulnerabilities. http://dx.doi.org/10.1101/534529.

O’Keefe C, McDevitt MA, Maciejewski JP. 2010. Copy neutral loss of heterozygosity: a novel chromosomal lesion in myeloid malignancies. Blood 115: 2731–2739.

Ried T, Hu Y, Difilippantonio MJ, Michael Ghadimi B, Grade M, Camps J. 2012. The consequences of chromosomal aneuploidy on the transcriptome of cancer cells. Biochim Biophys Acta 1819: 784.

Schwartz S, Papenhausen P. 2017. Significance of Copy-Neutral Loss of Heterozygosity Detected in Oncology Samples: Insights and Mechanisms. Cancer Genet 214-215: 40.

Stirewalt DL, Pogosova-Agadjanyan EL, Tsuchiya K, Joaquin J, Meshinchi S. 2014. Copy-neutral loss of heterozygosity is prevalent and a late event in the pathogenesis of FLT3/ITD AML. Blood Cancer J 4: e208–e208.

Suzuki H, Aoki K, Chiba K, Sato Y, Shiozawa Y, Shiraishi Y, Shimamura T, Niida A, Motomura K, Ohka F, et al. 2015. Mutational landscape and clonal architecture in grade II and III gliomas. Nat Genet 47: 458.

Taylor AM, Shih J, Ha G, Gao GF, Zhang X, Berger AC, Schumacher SE, Wang C, Hu H, Liu J, et al. 2018. Genomic and Functional Approaches to Understanding Cancer Aneuploidy. Cancer Cell 33: 676–689.e3.

van Jaarsveld RH, Kops GJPL. 2016. Difference Makers: Chromosomal Instability versus Aneuploidy in Cancer. Trends Cancer Res 2: 561–571.

Vattathil S, Scheet P. 2013. Haplotype-based profiling of subtle allelic imbalance with SNP arrays. Genome Res 23: 152–158.

Zack TI, Schumacher SE, Carter SL, Cherniack AD, Saksena G, Tabak B, Lawrence MS, Zhsng C-Z, Wala J, Mermel CH, et al. 2013. Pan-cancer patterns of somatic copy number alteration. Nat Genet 45: 1134–1140.

